# Genome scale identification of new genes using saturated reporter transposon mutagenesis

**DOI:** 10.1101/2024.09.06.611592

**Authors:** Emily C. A. Goodall, Freya Hodges, Weine Kok, Budi Permana, Thom Cuddihy, Zihao Yang, Nicole Kahler, Kenneth Shires, Karthik Pullela, Von Vergel L. Torres, Jessica L. Rooke, Antoine Delhaye, Jean-François Collet, Jack A. Bryant, Brian M. Forde, Matthew Hemm, Ian R. Henderson

## Abstract

Small or overlapping genes are prevalent across all domains of life but are often overlooked for annotation and function because of challenges in their detection. The advent of high-density mutagenesis and data-mining studies suggest the existence of further coding potential within bacterial genomes. To overcome limitations in existing protein detection methods, we applied a genetics-based approach. We combined transposon insertion sequencing with a translation reporter to identify translated open reading frames throughout the genome at scale, independent of genome annotation. We applied our method to the well-characterised species *Escherichia coli* and identified ∼200 putative novel protein coding sequences (CDS). These are mostly short CDSs (<50 amino acids) and in some cases highly conserved. We validated the expression of selected CDSs demonstrating the utility of this approach. Despite the extensive study of *E. coli*, this method revealed proteins that have not been described previously, including proteins that are conserved and neighbour functionally important genes, suggesting significant functional roles of these small proteins. We present this as a complementary method to whole cell proteomics and ribosome trapping for condition-dependent identification of protein CDSs, and as a high-throughput method for testing conditional gene expression. We anticipate this technique will be a starting point for future high-throughput genetics investigations to determine the existence of unannotated genes in multiple bacterial species.

## Introduction

The last two decades have seen an substantial increase in sequenced bacterial genomes, coupled with an increase in the associated pan-genome of any given organism. However, in recent years there has been a growing body of work to suggest that our annotation of bacterial genomes is incomplete. First, smaller proteins have been historically overlooked in all domains of life ^1–4^. Early definitions of a gene or coding sequence applied a size cut-off for annotation of a gene, ranging from 50-100 codons. However, several translated open reading frames (ORFs) as small as two or three codons have been identified in *Escherichia coli* and other species ^5–7^. A small protein, as defined by Storz and colleagues, is directly translated from an ORF (as opposed to processed from a larger protein), and composed of fewer than 50 amino acids ^4^. Small proteins (also referred to as “short ORF-encoded proteins” (SEPs), “microproteins” or “sproteins”) have been identified across the bacterial kingdom (reviewed extensively elsewhere ^8–10^), with roles in stress response (e.g. Prli42, TisB ^11–13^), cell division (e.g. MciZ, SidA, Blr ^14–17^), metabolism (e.g. SgrT, MgtS, MntS ^18,19^), respiration (e.g. CcoM, CydX ^20,21^), signal transduction (e.g. Sda, MgrB ^22,23^), modulating antibiotic resistance (e.g. Blr, AcrZ, Prli53 ^6,24^), defence and toxin/antitoxin systems (PepA1, TisB, Fst ^12^), or mediating host-interaction (e.g. rio3 ^25^) and survival inside host cells (KdpF ^26^). Given their functional importance, a number of studies have sought to identify these previously overlooked small proteins at scale ^27^, but this has not been without its challenges. Small proteins are harder to predict bioinformatically as the short sequence length makes it difficult to distinguish true coding sequences from random in-frame sequences between a start and stop codons ^28^. Furthermore, they are difficult to predict through sequence conservation analysis as some small proteins have species-specific function and therefore are genus or lineage-restricted.

Another class of overlooked genes are nested genes. Nested genes are defined as those encoded within another gene. They can be encoded within the same orientation as the primary coding sequence or anti-sense to the primary transcript. A subset of nested genes are isoforms^29^, which are encoded from an alternate start codon within the primary transcript, such as *cheA*(L) and *cheA*(S) in *E. coli* ^30^, or the three isoforms of translation initiation factor 2 encoded by *infB* ^31,32^. Nested genes are prevalent in viruses and eukaryotes due to their compact genomes or coding complexity, respectively, but considered rare anomalies in bacterial genomes ^33^. As such, many automated bacterial genome annotation tools discard putative coding sequences that are within larger genes. However, well documented examples of (non-isoform) nested genes include the *Shigella* enterotoxin (ShET) 1 gene encoded antisense to the Pic autotransporter ^34^, both virulence factors of diarrhoeagenic *E. coli* isolates, and *comS* encoded in the same sense but out of frame (OOF) to *srfA* in *Bacillus subtilis,* both of which have complementary but independent functions in natural competency and surfactin synthesis and secretion, respectively ^35^. More recent discoveries of overlapping genes (*olg1* and *olg2* respectively) as large as 957 and 1728 nucleotides have been demonstrated in *Pseudomonas aeruginosa* ^36^, suggesting overlapping protein coding is not limited to just small proteins.

Due to the limitations outlined above, high-throughput methods to identify protein coding sequences are needed to fully elucidate the genetics and biology of a given organism. Specialist methods that trap mRNA-bound ribosomes (Ribo-seq) have been developed to identify translated mRNA and therefore protein coding sequences ^29,37,38^. Although these studies highlight the remarkable extent to which small proteins have been overlooked, the data can however pose challenges for interpretation ^39^. Ribo-seq methods have been further optimised through the use of antibiotics that trap translating ribosomes at specific positions along translated mRNA^40^; one example termed ‘Ribo-Ret’ uses retapamulin, which traps initiating ribosomes resulting in enrichment of ribosomes localised near the initiating start codon. Such adaptations improve the resolution of the output data, but pose other challenges such as differentiating true coding sequence from ‘pervasive translation’ ^41^. Small proteins are also challenging to detect via mass spectrometry (discussed in detail by Ahrens *et al.* ^42^). They might be transiently expressed, expressed only under specific conditions ^43^, low in abundance, have few sites for trypsin digestion, or highly hydrophobic membrane-associated proteins. Taken together, it is clear that complementary methods are needed if we are to fully realise the coding potential of any given organism.

Translation reporters are one tool for identification of protein coding sequences. Such assays involve the fusion of the C-terminus of the target protein to the N-terminus of an enzyme or protein with a measurable read out, such as *β*-galactosidase, alkaline phosphatase or a fluorescent protein. Most translation reporter assays have been applied to defined target sequences, often in isolation. However, an early transposon-based random mutagenesis approach introduced the *phoA* gene, encoding alkaline phosphatase, into a Tn*5* transposon derivative ‘Tn*phoA*’. The transposon was designed such that the alkaline phosphatase, which is only functional in the periplasm, lacked its signal peptide sequence and therefore would only be functional if fused to sequences that promote its export ^44^, thereby identifying secreted proteins and periplasmically located portions of integral membrane proteins. This approach pre-dated the high-throughput amplicon-sequencing technology available today but demonstrates the utility of a transposon-based reporter system. Here, we combined a mini-Tn*5* translation reporter with ultra-dense transposon mutagenesis and sequencing to query the *E. coli* K-12 genome for new protein coding sequences independently of existing genome annotation at scale. By using this approach, we identified previously uncharacterised protein coding genes. We also document their conservation among >200,000 *E. coli* genomes. The capacity to discern an organism’s coding potential in a single experiment promises to be a powerful strategy for unveiling genotype-phenotype relationships.

## Results

### Construction of a reporter-transposon library

In our previous research^45^, the resolution afforded from ultra-dense transposon mutagenesis highlighted the presence of newly annotated small genes that contribute to cell fitness. We hypothesised that there might be additional as yet unannotated coding sequences in the *E. coli* genome, and that a transposon mutagenesis approach could be leveraged for their detection. A reporter transposon was constructed by introducing a kanamycin resistance gene upstream of the chloramphenicol resistance cassette of the transposon. However, the transposon was designed such that the aminoglycoside-3’-phospotransferase (*aph*) gene that confers kanamycin resistance was introduced immediately after the mini-Tn5 inverted repeat without a promoter, ribosome binding site (RBS) or start codon (Fig. 1). Expression of the *aph* gene is dependent upon in-frame insertion of the transposon into an actively translated gene. We first confirmed that the intact antibiotic selection cassettes did not confer any cross resistance (S. Fig. 1A). We next verified that the *aph* gene could confer kanamycin resistance with an N-terminal fusion of the transposon inverted repeat at the 5’ end of the *aph* gene. The transposon was cloned in-frame into LacZα in a pUC19 vector such that expression of *aph* was dependent upon Plac and the start codon of LacZα. The LacZα-Tn fusion was sufficient to confer kanamycin resistance and this resistance was lost if the LacZα start codon is mutated from ATG to AGG (S. Fig. 1B-C). Having confirmed the functionality of the transposon, we introduced the transposon via transposition into the *E. coli* K-12 strain BW25113 genome at random aiming for 1 transposon insertion event per cell. Transposon mutants were selected on LB agar plates supplemented with chloramphenicol and were pooled to form the library. The transposon-genomic DNA (gDNA) junctions were sequenced to identify the transposon insertion site. We designed our sequencing primers to sequence from the kanamycin resistance gene into the gDNA upstream to directly identify CDS translation fusion sites (S. Fig. 2). We sequenced two aliquots of the library and obtained 4.2 M and 2.7 M sequencing reads per replicate respectively. Comparison of the insertion density per gene between technical replicates showed sequencing the transposon junction was reproducible between replicates with a Pearson correlation coefficient of 0.99 (S. Fig. 3A), therefore we pooled the replicate data resulting in a total of ∼6.9 M mapped reads. Sub-sampling of our transposon junction sequences confirmed we had obtained sufficient sequencing for full coverage of our library (S. Fig. 3B). We identified 185,709 and 184,405 unique insertions for each orientation of the transposon respectively representing a total of 370,114 translational reporter mutants. When we further delineated the data by reading frame there was an equivalent number of mutants for each frame (S. Fig. 3C), distributed evenly throughout the genome (S. Fig.3D).

**Figure 1.**
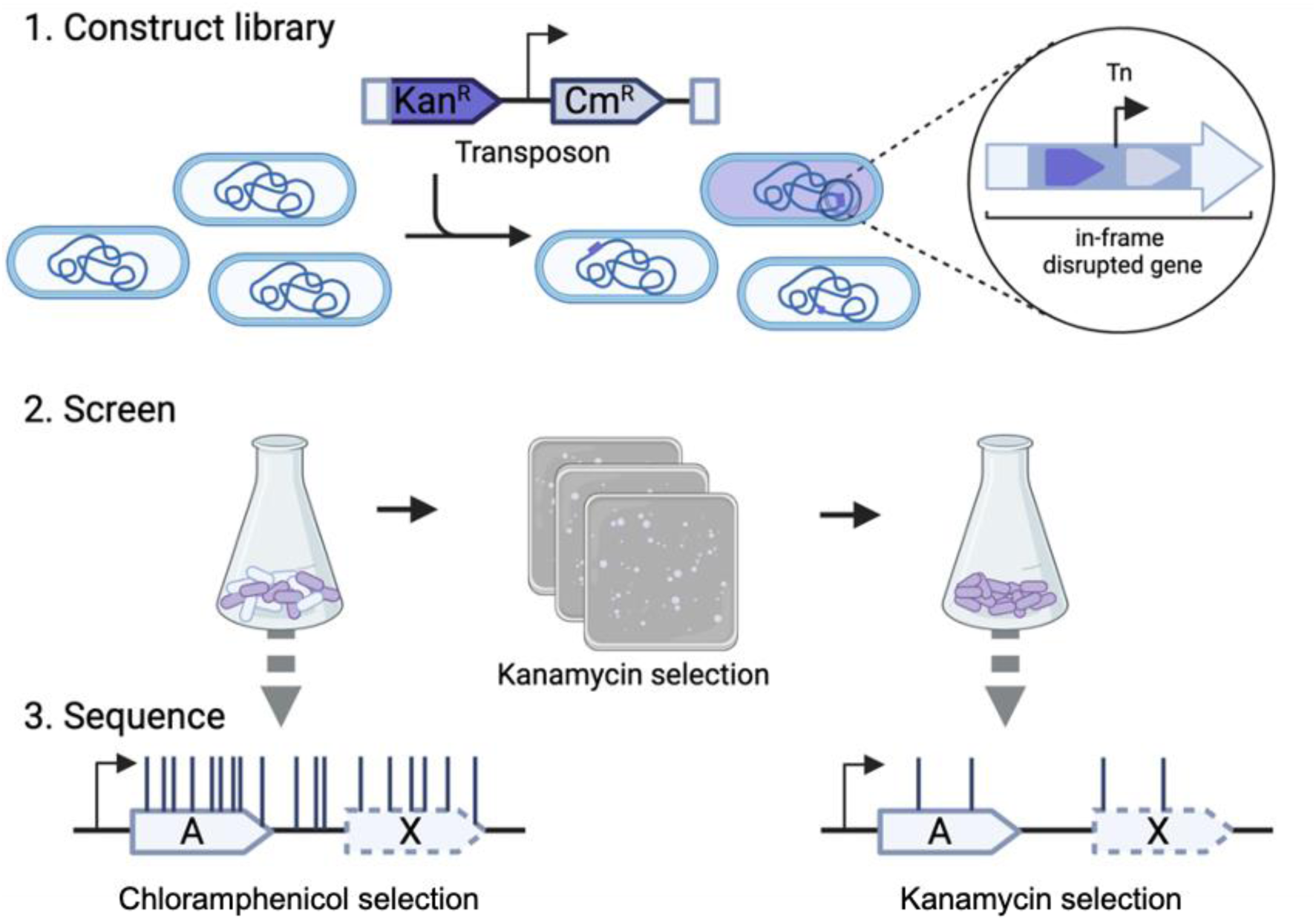
Method overview. (1) Construction of a transposon mutant library via introduction of a transposon with a dual selection mechanism. Successful transformants are isolated via selection on LB agar supplemented with chloramphenicol. (2) Screening of the transposon library on LB agar supplemented with kanamycin enriches for mutants that contain a transposon inserted in-frame in an expressed protein coding sequence. (3) Sequencing of the input and output transposon mutant pools reveals fusion mutants that resulted in the expression of the kanamycin resistance cassette, and therefore identifies protein coding sequences.

### Identification of putative new genes

To identify expressed protein coding sequences, we repeated selection of the library on agar plates supplemented with an inhibitory concentration of kanamycin, with the expectation that kanamycin resistance will only be conferred if the transposon has inserted in frame into an expressed protein coding sequence. For selection of kanamycin resistant mutants, 25 µg/ml of kanamycin was identified as the minimum concentration of kanamycin that inhibits growth of *E. coli* BW25113 on supplemented LB agar plates while selecting for transposon mutants (S. Fig. 4A-B). We chose to screen the library at this concentration to minimise the selection stress and maximise the number of translation-fusion mutants recovered. The library was screened twice, on separate days. We sequenced the transposon-gDNA junctions as before and recovered >150,000 reads per sample, which mapped to ∼8,000 unique insertion sites per replicate. The number of recovered unique insertions was lower than the number of fusion mutants we expected to recover. This may be a result of disrupted mRNA stability when endogenous CDSs are disrupted mid-sequence: indeed, review of the relative position of insertion sites found they were predominantly within the extreme 5’ or 3’ of each CDS (S. Fig. 4D). The biological replicates of mutants selected on kanamycin plates had a strong Pearson correlation coefficient (*r* = 0.81) between identified insertion sites (S. Fig. 4C). To account for the possibility that individual mutants might grow if they acquire spontaneous kanamycin resistance, we filtered the data to include hits that were identified in both independent screens, resulting in a final number of 6,219 transposon insertion sites. We then reviewed the functionality of the screen: we split the insertion data by reading frame and confirmed the bp level of resolution for identification of translated reading frames. For example, within the *tig-clpP-clpX-lon-hupB* operon the transposon insertions were each consistent with the appropriate reading frame for translation fusion with the kan^R^ *aph* gene (Fig. 2), demonstrating the functionality of the screen. Moreover, this level of resolution can reveal putative CDSs that do not coincide with annotated genes, for example three unannotated putative novel CDSs were identified within the *tig* operon (Fig. 2).

**Figure 2.**
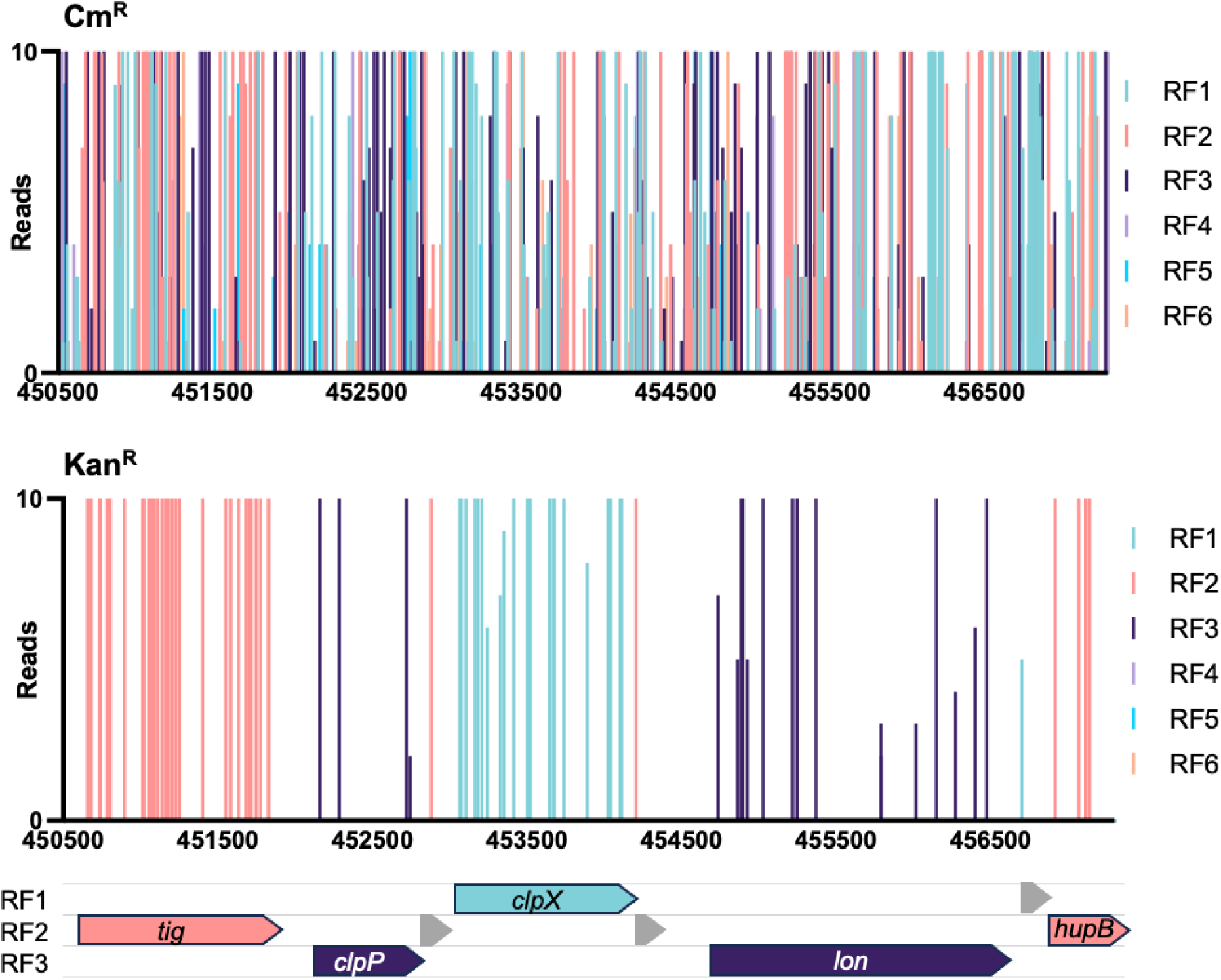
The bp resolution of the translation reporter reveals the reading frame of protein coding sequences. The initial construction of the transposon library was selected on agar plates supplemented with chloramphenicol (top panel). The vertical bars represent both the frequency and position of identified transposon insertion sites. The data are coloured according to the reading frame (RF) at the site of insertion. A secondary selection on agar plates supplemented with kanamycin selected for mutants where the transposon is inserted in frame of a protein coding sequence. The data are coloured according to the reading frame at the point of insertion and are consistent with the annotated genes (bottom panel). Annotated genes are indicated by black arrows while putative new protein coding genes are indicated by grey arrows.

Of the conservative 6,219 insertion sites, 5,754 insertions (92.52%) were in-frame within 1,139 annotated genes, 272 insertions (4.37%) were within annotated genes but out of frame and 193 insertions (3.10%) were intergenic (between annotated protein coding sequences; Fig. 3A). We used the well curated GenBank sequence (accession CP009273) as a reference for protein coding genes, however, to account for more recently annotated genes we also cross referenced our data against the RefSeq annotation (annotated by the NCBI prokaryotic genome annotation pipeline [PGAP], accession NZ_CP009273, annotated 24-APR-2023). This led to the positive identification of an additional 7 genes, 5 of which (*mgtT*, *pssL*, *ynfU*, *yqgH* and *ysgD*) were identified in a targeted Ribo-Ret screen in *E. coli* K-12 strain MG1655, further validating the utility of our approach to identify small coding sequences ^38^. Of the remaining two, *rseD* and *nadS*, *rseD* has been previously validated by chromosomal integration of a *lacZ* translation reporter ^46^. While *nadS* (RS26030), although not currently listed in Ecocyc, is annotated in the PGAP reference genome as a hypothetical protein, our data presents the first lab-based evidence for its expression and suggests it is expressed under normal laboratory growth conditions. Overall, the revised total of known genes that were identified in the screen was 1,146. We cross-referenced our data against other whole-cell protein-identification datasets, including a proteomic mass-spectrometry dataset and a Ribo-Ret dataset ^29,47^, chosen for using a comparable strain (*E. coli* BW25113 or derivative) and growth condition (LB +/- supplements; S. Table 1). The largest cohort of proteins were shared by all three methods (S. Fig. 5A), however there were clearly some method-dependent differences in the proteins identified (discussed extensively elsewhere ^39,48,49^). Transposon mutagenesis screens are inherently limited by their inability to screen regions of the genome that cannot viably be disrupted, such as essential genes. However, we were able to detect some essential genes, such as within the *mur* operon (S. Fig. 5B), where the transposon is in-frame within the extreme 5’ or 3’ end of the protein CDS, the former due to translational read-out from the chloramphenicol resistance marker maintaining downstream expression ^45^. Thus, detecting essential proteins is possible with transposon mutagenesis, but requires saturating mutagenesis.

**Figure 3.**
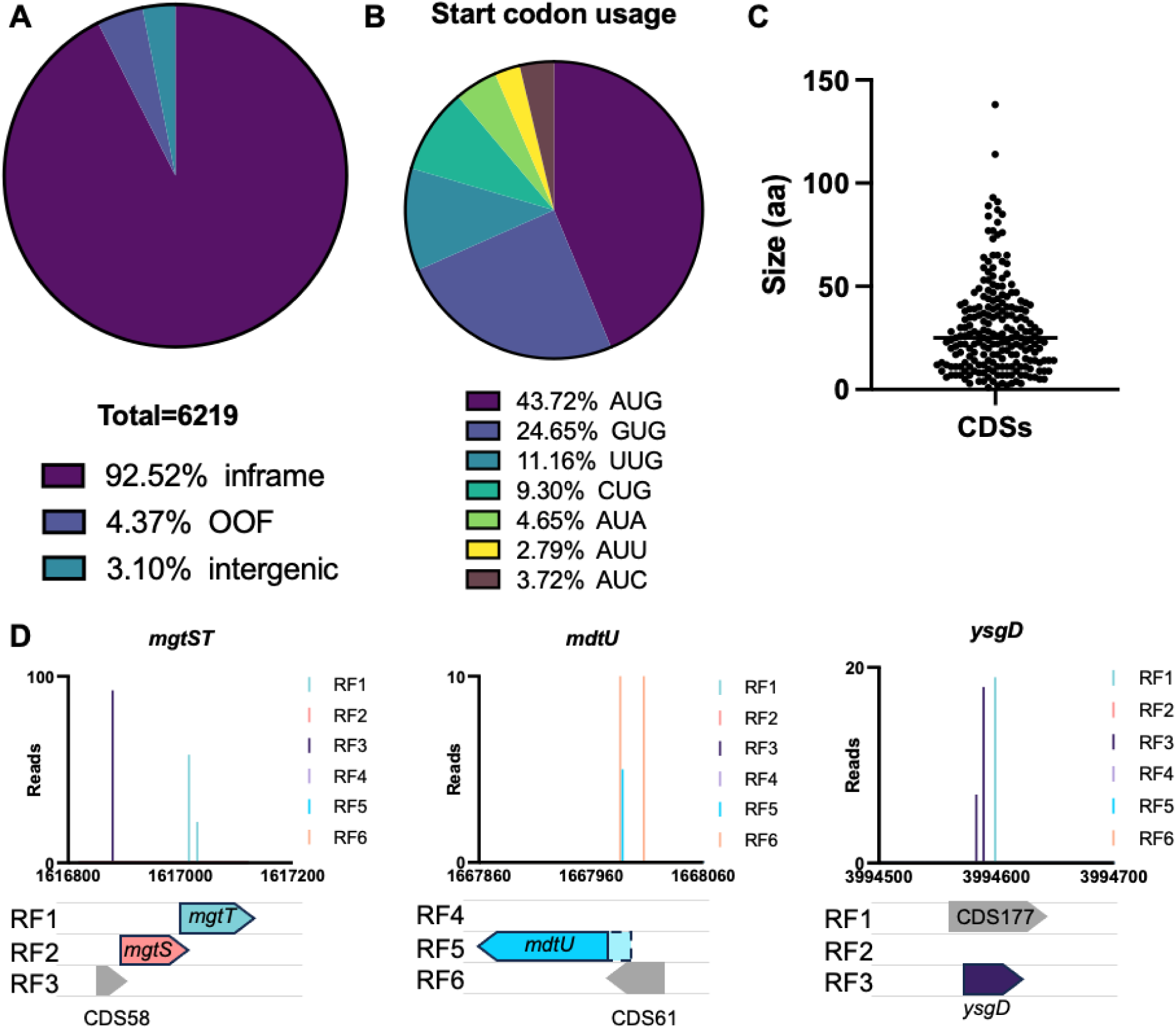
Identification of protein coding sequences. (A) The identified transposon-insertion site (and therefore translation fusion junction) was mapped to the reference genome and cross-references with annotation information. 92.52% of insertions were within annotated genes and ‘inframe’ consistent with the reading frame of the annotated gene. 4.37% of insertions were within annotated genes but in a different reading frame to the annotated gene (“out of frame [OOF]) and 3.10% of insertions were not within annotated protein coding sequences. (B)(i) The start codon frequency of 215 putative CDSs, and (ii) the CDS sizes in amino acids, with the median shown by a horizontal line. (C) Small CDSs identified by reporter TIS neighbouring small genes identified by Ribo-Ret^38^. The transposon insertion sites are those identified following selection with kanamycin (representing translation-fusion events) and are coloured according to the reading frame (RF) at the site of translation-fusion. Putative new genes are shown in grey.

To identify putative new genes, we reviewed the remaining translation-fusion sites that were either within genes but out-of-frame, suggestive of a nested gene, or intergenic and therefore candidate new coding sequences. While the translation-fusion data can reveal open reading frames that are translated, it cannot identify which codon is initiating translation. Although the most common start codon in *E. coli* is AUG, one study found 47 (out of 64) codons are capable of initiating translation, albeit at varying strengths ^50^. To be conservative we focused on the strongest 7 codons that can initiate translation: AUG, GUG, UUG, CUG, AUA, AUU and AUC, which all displayed expression levels >100-fold higher than the control cells ^50^. We reviewed both the nested ‘out-of-frame’ and intergenic reading frames identified in our translation screen for these start codons and identified a total of 215 unique putative protein coding sequences (S. Table 2); 125 were nested gene candidates while 90 were intergenic. Most annotated genes containing a predicted nested gene had only 1 internal nested gene (S. Fig. 6), however, there were a handful with multiple predicted nested genes. Although multiple nested genes within a parent gene has been previously documented in *E. coli,* such as within *waaL* which contains 3 nested genes ^38^, another possible explanation for multiple nested candidates within the TIS data is that these arise from ribosome slippage during translation of the primary transcript, as *E. coli* K-12 strains are prone to higher rates of ribosome slippage than other *E. coli* lineages ^51^, and the genes with ≥5 nested genes are some of the most highly expressed proteins ^47^.

The predominant predicted start codon in the 215 candidate CDSs was AUG (43.72%) followed by GUG (24.65%) and UUG (11.16%) (Fig. 3B). Although the initial aim of this screen was to identify protein coding sequences irrespective of genome annotation, we observed that most of the new CDSs were small proteins <50 amino acids (aa; median 25 aa, smallest 1 aa, largest 138 aa; Fig. 3C). We did not apply a lower limit threshold for the size of putative CDS as (1) translation of 2 codon sequences is known for *E. coli*, although these sequences often represent translation-control mechanisms rather than a functional protein and (2) the calculated sizes are derived from an estimated start codon and therefore might not be accurate. The finding that most putative new CDSs were <50 aa is not entirely surprising and is consistent with the hypothesis that small proteins are prevalent yet have been systematically overlooked during bacterial genome annotation. Surprisingly, comparison of the candidate 215 CDSs with the 68 candidate small proteins identified by Ribo-seq methods by Weaver *et al*. ^38^ revealed only 9 genes that were identified by both methods. Of note, many of the small proteins identified uniquely by Ribo-Ret appear recalcitrant to transposition, precluding them from our screen. This raises the possibility that small genes have specific regulatory mechanisms that limit their disruption or represent an overlooked class of essential genes. However, many of the small genes detected using both methods have additional neighbouring putative small genes within our data, suggesting organised transcriptional units of small genes. For example, our data is in agreement with the Ribo-Ret data for the identification of small genes *mgtT*, *mdtU* and *ysgD*, but we also detected translation in alternate reading frames neighbouring these genes (Fig. 3D). For *mgtT* and *mdtU*, we identified translation upstream of these genes, which coincided with putative CDSs with a AUG start codon (CDS58 and CDS61 respectively), yet these were not detected by Ribo-seq. Moreover, within *mdtU* we observed translation earlier within the reading frame than the given start codon for *mdtU*, suggesting an alternate start codon further upstream. For *ysgD*, we detected translation both within this reading frame and within an overlapping reading frame, consistent with CDS177, which also has a putative start codon (AUC). Interestingly, Weaver *et al.* ^38^ observed translation extending beyond the stop codon of *ysgD*, which would be consistent with expression of CDS177, but the absence of an AUG, GUG or UUG start codon filtered CDS177 from their analysis.

### Validation of new genes

To assess the validity of our method, we selected 17 putative new CDSs identified by this method for verification by western blot analysis. We chose candidates with a cross-section of genomic neighbourhoods (start-, mid- and end-operon), in addition to some nested genes for proof of principle, and focused on these 17 because of their gene-neighbourhood with some well-known and extensively studied genes (Fig. 4A). Of the 17 putative CDSs, 12 are intergenic and candidate operonic genes, while the remaining 5 are nested genes but in a different reading frame to the ‘parent’ annotated gene; 1 of the nested genes (within *damX*) was only identified in one of the kanamycin-selection replicate datasets, but because of its unusual transposon insertion profile we included it for follow-up investigation. Using lambda-red mediated recombination, a dual epitope “sequential peptide affinity” (SPA) tag, comprising a calmodulin binding peptide and 3×FLAG tags, was inserted into the genome immediately upstream of the native stop codon. Constructs were checked by PCR (S. Fig. 10) and confirmed by Sanger sequencing. This approach has been demonstrated previously for the detection of small proteins ^52,53^, as the large size of the tag (∼8 kDa) can enable detection of proteins as small as 1 kDa. Strains were grown in LB broth to both exponential and stationary phase, cells were harvested and the whole cell lysates probed with anti-FLAG antibody for the detection of endogenously expressed small proteins. We detected expression for 6/12 intergenic CDS, and 3/5 of the nested CDSs (Fig. 4B). Small proteins in *E. coli* vary in their detection levels by western blot analysis ^52^, and there were visible differences in the levels of protein detected here. CDS58, encoded upstream of *mgtST*, had the weakest signal suggesting this protein is not stable or is present at very low levels. We observed that the number of reads for each small protein did not correlate with the strength of western blot detection, for example, CDS58, which is the faintest on a western blot, had 93 reads, whereas CDS119 and CDS147 were very prominent by western yet only represented by 9 and 12 reads respectively. As such, we did not set a minimum read threshold for CDS detection provided the CDS was detected in two independent kanamycin-selection screens. It’s important to note that western blot detection of the C-terminal FLAG tag can validate endogenous translation of a given reading frame, but cannot confirm a predicted start codon for initiating translation. For example, we detected the expression of tagged CDS66, however, the migration of this protein was indicative of a larger protein than expected. We confirmed by Sanger sequencing that the correct reading frame was targeted, and the SPA tag is out-of-frame with the *ompC* start codon. Several possible explanations are (1) a homomultimeric complex that is not reduced by the denaturing conditions when separating the proteins, (2) translation is initiated from the *ompC* start codon and a +1 frameshift enables expression of the SPA tag, however the predicted size of this construct (with tag) is ∼34.5 kDa, (3) the amino acid sequence gives rise to aberrant migration such that the observed size is not consistent with the predicted size, as has been observed for Antigen 43 ^86^, or (4) the protein undergoes some form of posttranslational modification that gives rise to aberrant migration on SDS-PAGE, as has been observed upon lipidation of the *E. coli* CexE protein ^87^.

**Figure 4.**
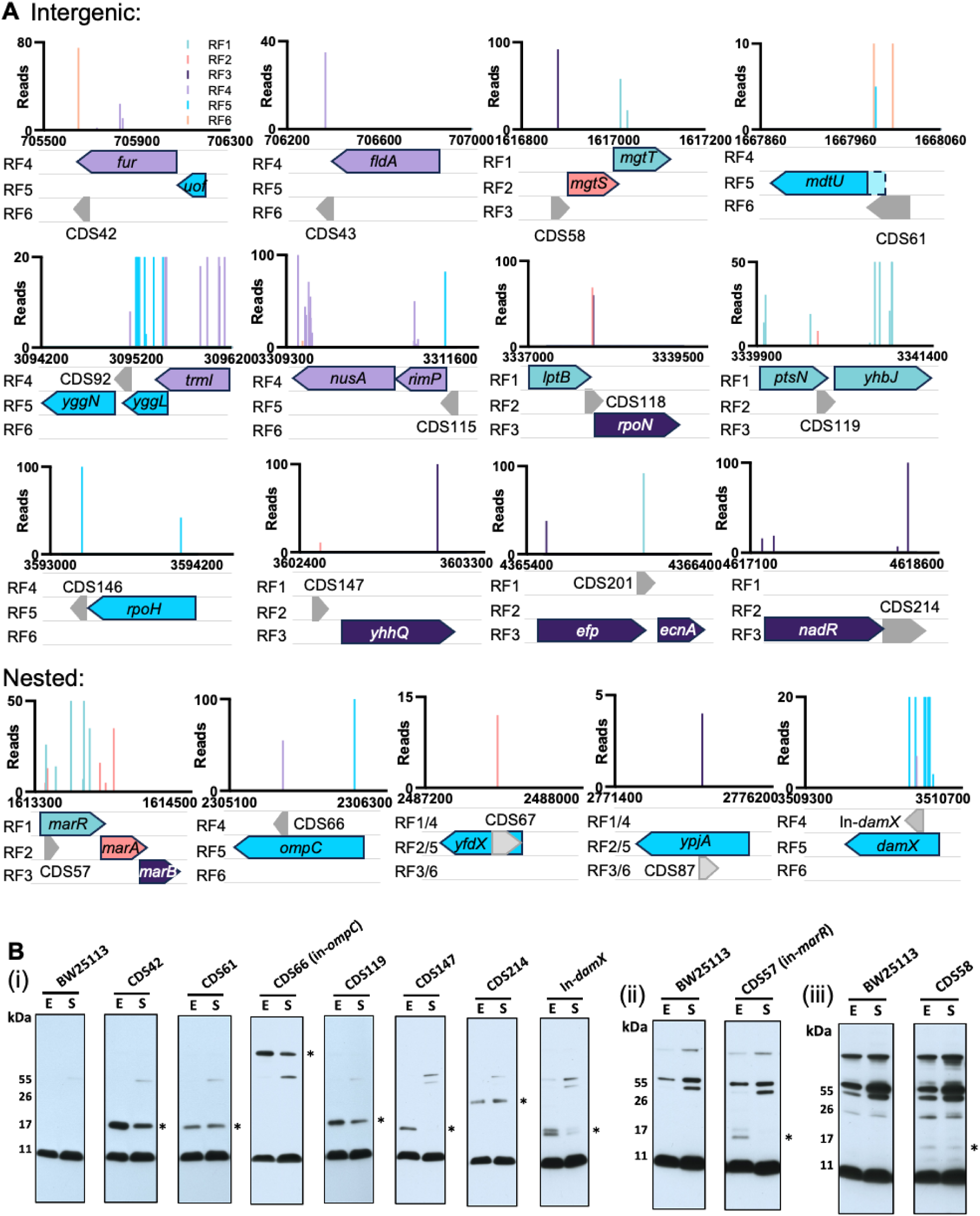
Validation of new proteins. (A) The genetic neighbourhoods of putative protein coding sequences (CDS) identified by reporter transposon-insertion sequencing selected for validation. Putative CDSs are shown in grey. (B) Representative western blots (of 3 repeats): whole cell lysates probed with anti-FLAG antibody, detected proteins are indicated by *. BW=BW25113 control (untagged); E = Exponential phase; S = Stationary phase. Blots are separated into three panels according to the exposure time used for protein detection (i) short (ii) medium (iii) overnight.

Of the CDSs that we did not detect, it’s possible that they are only transiently expressed, are susceptible to rapid degradation, are expressed at levels below the threshold for western blot detection, or are destabilised by C-terminal location of the tag. Indeed, a study validating small proteins identified by Ribo-seq methods found that some could not be detected by western blot analysis until the addition of a ClpP protease inhibitor, bortezomib ^41^. Another possibility is that some of these proteins might be secreted, and therefore missed during our cell harvesting steps prior to lysing the cells for western blot analysis. However, none of the putative CDSs contained any predicted signal sequences ^54^.

We next predicted the secondary and tertiary structures of the small proteins using PSIPRED and AlphaFold2, including transmembrane (TM) domain prediction using MEMSAT-SVM. CDS42, CDS43, CDS58, CDS61, CDS146, CDS201 and CDS57 were all too short for analysis by PSIPRED (minimum size is 30 residues). All proteins that passed the size threshold for PSIPRED and MEMSAT analysis were predicted to contain a single TM helix, with 4 of these (CDS118, 119, 147 and 87) predicted to be pore-lining (S. Fig. 7). The AlphaFold2 predictions, generated by the ColabFold pipeline, largely corroborated the secondary structure predictions of PSIPRED, with many helical domains predicted for these small proteins (S. Fig. 8). Although the confidence metrics (pLDDT and pTM) of these predicted structures are low, this likely stems from the limited depth and diversity in the multiple sequence alignments (MSA) generated by the ColabFold pipeline as the AlphaFold2 system relies on deep and diverse alignments to generate high-confidence models ^55,56^, however, newly identified proteins may have small sequence families or few homologs in the ColabFoldDB database through which the MSA are generated ^55,56^. Although small proteins are notoriously challenging to classify, the data from three independent structural prediction tools suggest some of these small proteins may be small helical proteins and possibly membrane proteins. Previous studies have reported the prevalence of helical proteins in small protein screens ^53^, with functions ranging from toxin systems that promote persister cell formation, to accessory subunits of larger membrane protein complexes ^57,58^.

### Conservation of new genes

Finally, we evaluated the conservation of these genes. To investigate the prevalence and distribution of the 17 candidate CDSs among *E. coli* strains, we screened a collection of 227,873 globally distributed *E. coli* genomes. Using tblastn analysis, we identified that most of the CDS were conserved in most (>95%) *E. coli* genomes (Fig. 5); with the exceptions of CDS146 (92.8%), CDS66 (in-*ompC*; 89.8%), CDS57 (in-*marR*; 87.1%) and CDS87 (in-*ypjA*, 44.8%), which appear to show some lineage restriction and might suggest these are more recently acquired or evolved genes (S. Fig. 9). For 8 of the small genes, we identified putative homologs outside *E. coli* (S. Data 1).

**Figure 5.**
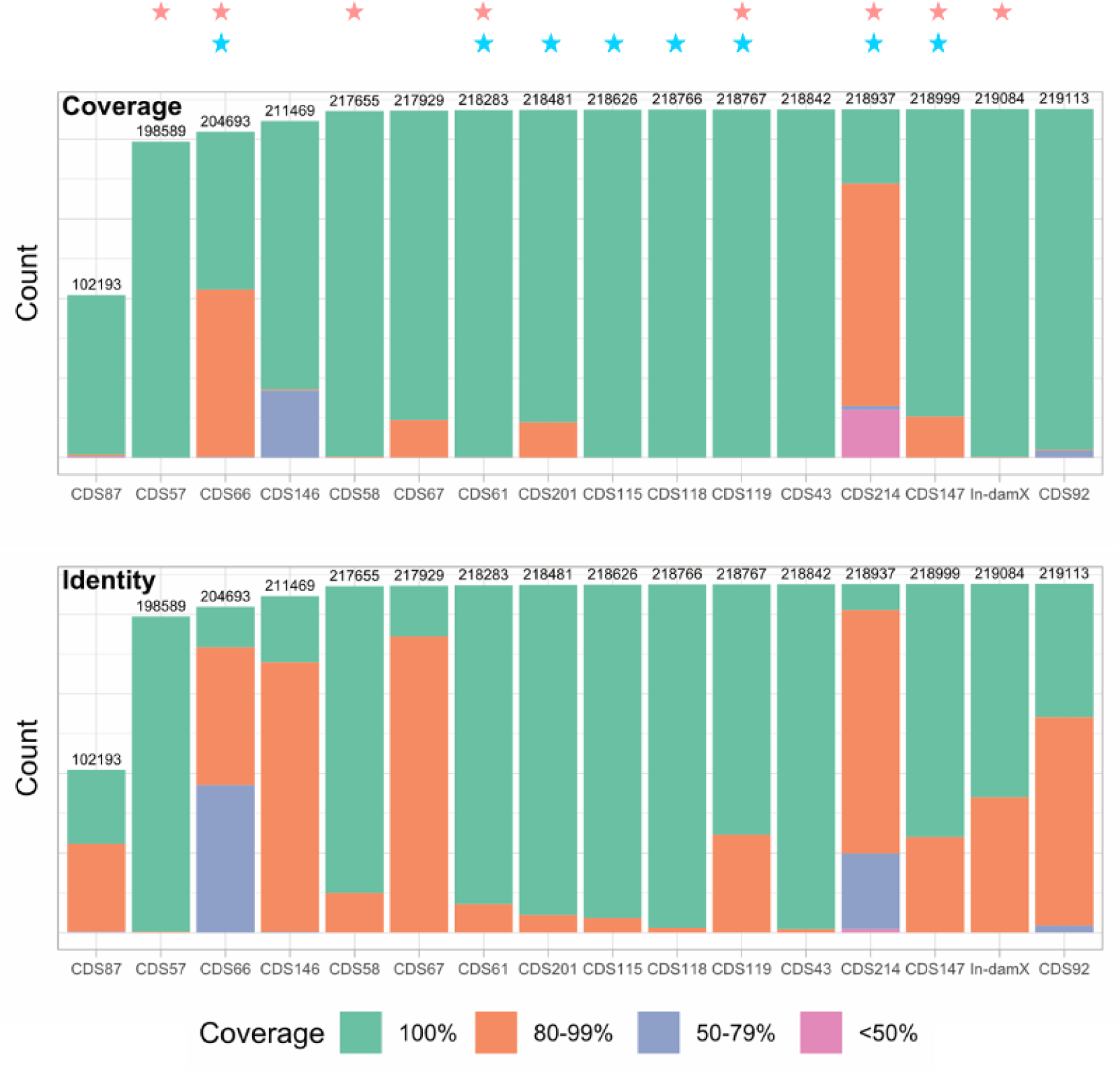
CDS carriage within the 227,873 genome database. tblastn results of CDS homologs within *E. coli* gnomes, coloured by % coverage and % identity accordingly. CDS validated by western blot analysis are indicated with a pink star (top). CDS with putative homologs outside of *E. coli* are indicated with a blue star (second track).

To establish whether the small genes are co-conserved with their gene neighbours, we reviewed whether the genetic synteny of the CDSs was preserved across the different *E. coli* isolates. A 4 kb window (2 kb up- and down-stream) surrounding each CDS was extracted from the *E. coli* BW25113 genome as a reference. The 4 kb region was compared by blastn to the corresponding *E. coli* database strains with identified homologs, to establish whether local gene neighbourhoods were maintained. Most CDS were conserved in ≥99% of isolates with at least one gene-neighbour indicating co-conservation (Fig. 6). For some genes, e.g. CDS43 neighbouring *fldA*, or CDS119 neighbouring *ptsN* and *yhbJ*, gene synteny was preserved along the full 4 kb region. Whereas CDS58 was more strongly co-conserved with the neighbouring *mgtST* operon than neighbouring *ydeE*, indicating the CDS58 is part of the *mgtST* operon, and in a subset of *E. coli* isolates (11.7%) this operon may be located elsewhere in the genome. While CDS115 was more strongly co-conserved with *rimP* suggesting a leader sequence for the *rimP-nusA* operon. Genes are often co-localised with functionally related genes and the level of conserved organisation might hint at the potential functions of these newly identified genes. Altogether the new small genes were co-conserved with genes with a range of biological functions, highlighting the wide degree of biological potential that may be overlooked by excluding small proteins.

**Figure 6.**
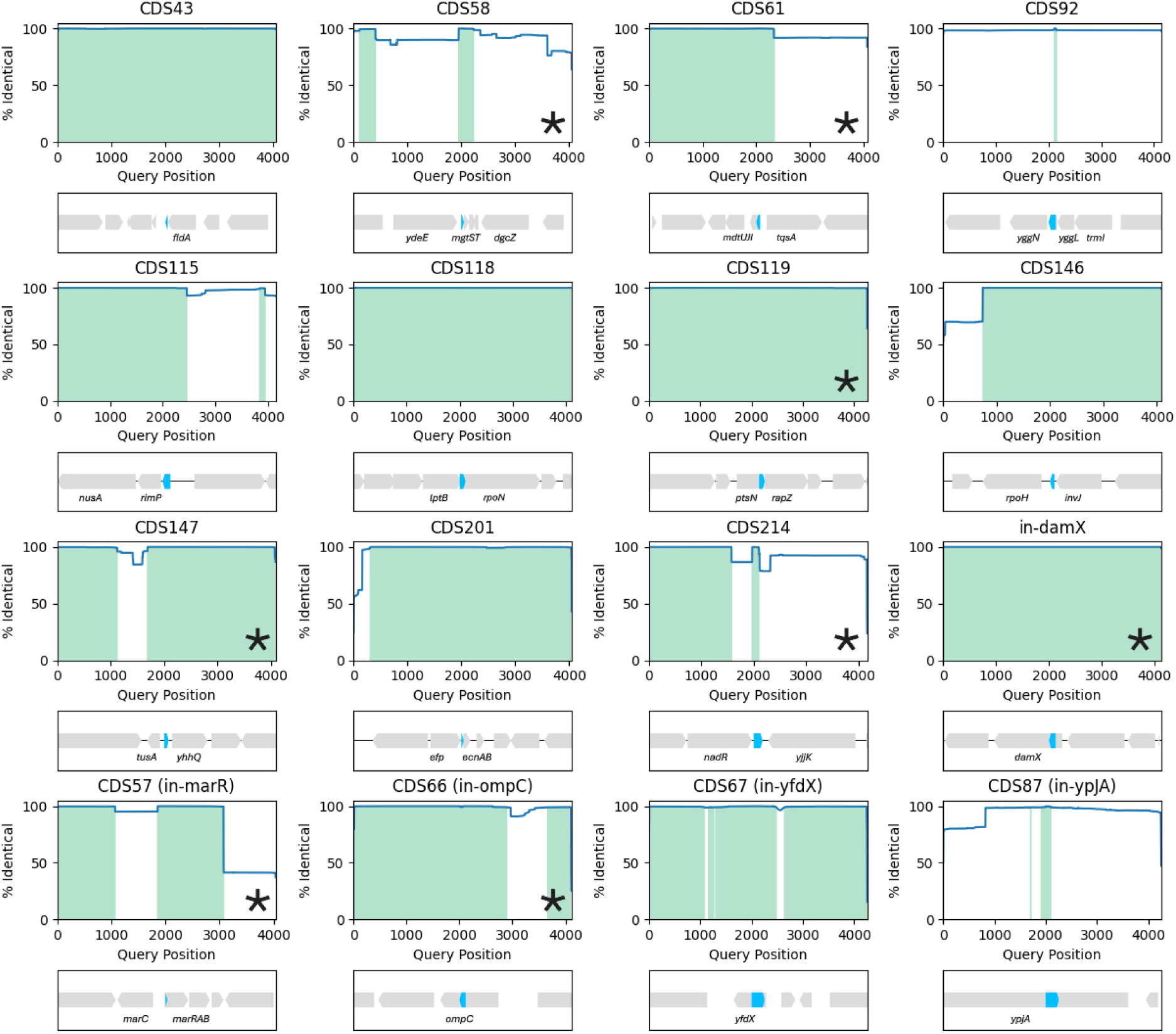
Nucleotide identity plots of gene neighbourhoods. Nucleotide sequence identity 2 kb up- and down-stream of each CDS from the *E. coli* BW25113 reference genome compared with the *E. coli* genomes database. Each plot shows the % of genomes (y-axis) that have coverage along the reference genome query sequence (x-axis) with the BW25113 reference sequence indicated below. Annotated genes are shown in grey, candidate new genes are highlighted in blue, the *mgrR* RNA (downstream *mgtST*) is coloured in green. CDS additionally detected by western blot are indicated with a star.

## Discussion

The *E. coli* genome has been highly scrutinised for the identification of novel small proteins ^9,41,52^, yet this method still revealed additional new proteins with the detection of >200 putative new coding sequences. We validated the endogenous expression of small proteins detected exclusively by this approach using western blot analysis, and confirmed detection of 9/17 of these CDSs, three of which are encoded within overlapping genes. Many of the newly identified genes were conserved throughout *E. coli* and often co-conserved with their genetic neighbours, with some genes predicted to have homologs in other genera, adding further support for their existence. Small proteins are increasingly being recognised for their diverse biological roles, which often includes the modulation and regulation of larger protein complexes ^59^. Our identification and detection of small proteins co-conserved with genes involved in electron transfer (*fldA*), magnesium transport (*mgtST* ^60^), ribosome maturation (*rimP* ^61^), spermidine efflux (*mdtUJI* ^62^), potassium transport (*ptsN* ^64^), regulatory RNA stability (*rapZ* ^65^), queuosine precursor transport (*yhhQ* ^66^) NAD+ biosynthesis (*nadR*) and tRNA methyltransferase (*trmI*), among others, suggests further functional contributions of small proteins and highlights the functional importance of small proteins that are still being overlooked. We present this as a high-throughput method complementary to Ribosome-trapping or mass spectrometry-based approaches for detecting expressed proteins.

Our gene neighbourhood conservation data and comparison with Ribo-seq data revealed conserved clusters of genes organised into transcriptional units of small genes, e.g. CDS58-*mgtST*, CDS61-*mdtUJI*, or CDS177-*ysgD*. The detection of additional genes not previously identified by Ribo-seq might be due to detection sensitivity (discussed below) but was in part due to our wider acceptance of putative start codons: where Weaver *et al*. used a conservative shortlist of AUG, GUG and CUG codons ^38^, we allowed for 7 putative start codons, which enabled the positive identification of CDS177 overlapping *ysgD*. However, these data are likely still an underestimate of the total proteome as earlier work using a GFP reporter has shown as many as 47 of the 64 codons (including one canonical stop codon) are able to initiate translation ^50^. Ribo-Ret analyses that allow for a wider selection of start codons observe more varied start codon usage ^29^, although this in part has been attributed to decreased ribosome selectivity due to increased 30S subunit availability. However, consistent with the hypothesis that non-canonical start codons can initiate translation, prior to filtering our data we did observe translation within intergenic regions with no obvious start codon. One recent study combining the analysis of stalled ribosomes at start and stop codons respectively identified almost 400 putative CDSs, however they attribute this high number in part to pervasive translation ^41^. While pervasive translation, described as the spurious translation of a non-functional coding sequences, could contribute to false positive translation-detection in our screen, we observed that the detection rate of expected functional translation-fusions was quite low (27.5%: the number of detected correct-RF insertions within annotated genes/the total number of correct-RF insertions for those genes) suggesting active translation alone is insufficient for positive translation-fusion detection, and that a degree of mRNA stability is required for successful aminoglycoside phosphotransferase expression.

It’s also important to note we don’t yet know the detection limits of western blot detection. For example, for two of the CDS we could not validate by western blot analysis, we did detect faint bands at the expected migration site with excessive blot contrast enhancement, although deemed these insufficient for validation and excluded them from our data. Additionally, there were CDS identified by the transposon-reporter that we could not validate by western blot, yet have conserved coding sequences (inclusive of start and stop codons) outside of *E. coli*, e.g. CDS115, CDS118, CDS201. It is possible that a transposon mutagenesis approach can offer an improved detection resolution as compared to detection by standard western blot, especially for small proteins that are too transiently or weakly expressed to be detected by standard western blot approaches without purification. As both the transposon translation reporter and SPA-tag detection by western blot inherently rely on the same principles of C-terminal tagging of endogenous proteins, an exciting option for future method-development here might be the introduction of a transposon-based SPA tag for whole-cell purification of small proteins.

One limitation of this method is that essential genes, or genes that are not easily disrupted, e.g. those bound by DNA-binding proteins, will be less well-represented within the transposon mutant pool, limiting their opportunities for detection. Secondly, this approach is inherently biased against the detection of small proteins, as shorter genes will also be represented by fewer translation-fusion mutants. Our data suggests that both of these limitations may at least in part be overcome using a mini-Tn*5* system that can achieve near saturated mutagenesis. A strength of this method is its bp level of resolution for identification of the precise reading frame where translation is occurring. However, as this method identifies functional translation fusion sites, there is always the possibility that an out-of-frame insertion is not evidence of a new gene, but evidence of ribosome slippage and a shift in the translation reading frame (programmed or otherwise).

Besides predicting new proteins, this method can also validate whether annotated genes are translated. One example of this in our data was within the ‘pseudogene’ *ilvG*. Curiously, our data suggests both that the N-terminal fragment of the pseudogene is still expressed, and that a second reading frame (+1) downstream is also expressed. We also detected translation within other unexpected genomic regions, including within a handful of repetitive extragenic palindromic (REP) elements. A phenomenon which has also been reported in another study measuring transcriptomics and proteomics in *E. coli* O157:H7 ^67^.

Although the method of coupling random mutagenesis with reporter systems is not novel ^44^, it is still surprisingly in its infancy when applied at scale, with only one other saturating mutagenesis reporter study (‘ProTinSeq’) published earlier this year in the model system *Mycoplasma pneumoniae* ^68^. As a method it is applicable to any organism that is genetically tractable and the scope of application is much broader than the system presented here. One obvious adaptation of this method is to replace the reporter marker with a non-lethal selection marker, such as a fluorescent protein, which, coupled with fluorescence activated cell sorting, would enable rapid *in vivo* screening for condition-dependent changes in expression. This would expand the capabilities for small protein detection as one often cited limitation is that small proteins might only be expressed transiently, at low levels, or under specific conditions. With many of these tools still in their relative infancy, and focusing on a limited range of growth conditions, it is likely there are still more small proteins to be discovered in the *E. coli* genome. In particular, nested genes have been gaining increasing interest and traction within the field ^69–71^; with data presented here adding another line of evidence in support of their wider existence.

Finally, this work highlights that a standardized approach to genome annotation is needed ^72,73^, and this has been recognised with the implementation of the NCBI Prokaryote Genome Annotation Pipeline applied to all newly submitted sequence data. However, we found that even the most “up to date” annotation of *E. coli* K-12 strain BW25113 (re-annotated 24-APR-2023) did not include many of the small proteins identified and validated in Ribo-seq experiments from 5 years ago within the same (or closely related) strains ^9^. As such, it’s worth noting that even for one of the best curated strains, *E. coli* K-12, many high-throughput functional genetics screens that rely on genome annotations for genotype-phenotype results are still overlooking small proteins.

## Materials and Methods

### Bacterial strains and culture conditions

*E. coli* K-12 strain BW25113 (*rrnB*_T14_ Δ*lacZ*_WJ16_ *hsdR514* Δ*araBAD*_AH33_ Δ*rhaBAD*_LD78_) was routinely used ^74^. Cell cultures were grown overnight in LB broth (Tryptone 10g/L, Yeast extract 5g/L, Sodium Chloride 10g/L) at 37°C supplemented with 100 µg/ml carbenicillin (Sigma) for plasmid selection, and 35 µg/ml chloramphenicol (Merck) or 25 µg/ml kanamycin (Sigma) for transposon selection.

### Cloning

The transposon sequence was constructed by stitch PCR using the commercially available EZ-Tn*5* Kan^R^ transposon (Lucigen), and a previously published Cm^R^ mini-Tn*5* transposon^45^ as a template and cloned into the pUC19 vector. The transposon was inserted between the *Hin*dIII and *Bam*HI restriction sites, resulting in the plasmid “pUC19-KC-Tn”, which contains the dual selection transposon inserted in-frame in the LacZa gene. Mutations of the LacZa start codon were introduced via site-directed mutagenesis. Briefly, the plasmid was amplified by PCR using phusion polymerase (NEB) and overlapping forward and reverse primers that contain the desired mutation. The vector template was digested by DpnI (NEB) following the supplier’s instructions. Newly amplified vector was transformed into competent DH5alpha cells (NEB) and correct mutations were confirmed by Sanger sequencing before transferring to *E. coli* strain BW25113.

### Construction of a transposon library

To make the transposome (transposon coupled with transposase), the pUC19-KC-Tn plasmid was linearised by restriction digest using *Nde*I (New England Biolabs) following the supplier’s instructions. The mini-Tn*5* transposon was amplified by PCR using primers with a phosphorylated 5’ end and containing the inverted repeat mosaic ends for efficient Tn5 transposase-mediated transposition. The transposon was amplified using the phusion polymerase (NEB) and the linearised plasmid as a template with the following conditions: 30 s 98°C; (10 s 98°C, 20 s 60°C, 1 min 72°C) x30; 5 min 72°C; 4°C hold. The PCR product was purified via gel extraction (Qiagen) following the supplier’s instructions. The Transposon was eluted in TE buffer (pH 8.0; ThermoFisher Scientific), quantified by Qubit dsDNA HS Assay kit (ThermoFisher Scientific) and adjusted to a final concentration of 100 ng/µl. To prepare the transposome, a ratio of 1:2:1 of 100 ng/µl transposon DNA in TE buffer, EZ-Tn5 Transposase (TNP92110; Gene Target Solutions) and 100% glycerol were incubated at RT for 1 h then frozen at -20°C for storage. The transposome was introduced into *E. coli* K-12 strain BW25113 by electrotransformation. Briefly, cells were grown to mid-exponential phase in 800 ml LB at 37°C with aeration, harvested by centrifugation at 2000 xg for 10 min at RT and washed 4x in 10% glycerol at the same volume. The final cell pellet was resuspended in 1 ml of 10% glycerol. 1 µl transposome was incubated with 100 µl cells before shocking at 25 kV. 1 ml LB was added per reaction, and cells were incubated a 37°C with shaking for 2 h for recovery. Successful transformants were selected on LB agar plates supplemented with 35 µg/ml chloramphenicol and incubated overnight at 37°C for 18 h. Colonies were scraped from plates to form the transposon library of pooled mutants and resuspended in 30% glycerol in LB for storage at -80°C. For selection of kanamycin resistant mutants, 25 µg/ml of kanamycin was identified as the minimum concentration of kanamycin that inhibits growth of *E. coli* BW25113 on supplemented LB agar plates (S. Fig. 5A) and was therefore selected as the concentration for selection of kanamycin-resistant isolates. For identification of mutants expressing the translation reporter, the transposon library was selected on LB agar plates supplemented with 25 µg/ml kanamycin, to achieve a density of ∼2,000 colonies per plate. A total of ∼1×10^7^ CFUs, of the input library were screened in total (per replicate), ensuring sufficient coverage of the library. Colonies were scraped from plates, resuspended in 30% glycerol-LB, pooled and stored at -80°C.

### Transposon sequencing

Genomic DNA was extracted, fragmented by ultrasonication to an average size of ∼250 bp and prepared for sequencing using the NEB Next Ultra I kit (E7370L, NEB), with some modifications. Following end repair, adapter ligation and a size-selection purification step with SPRI beads (Beckman Coulter) following the kit instructions, a PCR step was introduced to enrich for Transposon-gDNA junctions, using a forward primer specific for the transposon and a reverse primer specific for the adapter. 15 µl DNA fragments in sterile, nuclease free water, 25 µl Q5 polymerase, 2.5 µl 10 µM each primer and 5 µl nuclease free water to a final volume of 50 µl were mixed and amplified in a thermocycler with the following profile: 3 min 98°C; (15 s 98°C, 30 s 65°C, 30 s 72°C) x10; 1 min 72°C; hold 4°C. The sample was purified using SPRI beads at a ratio of 0.9:1 beads to sample. A second PCR step then prepared the transposon-gDNA junctions for sequencing using custom forward primers specific for the transposon, which incorporated a barcode for sample identification as well as staggering the start of the transposon sequence during sequencing. The reverse primers, NEBNext Multiplex Oligos for Illumina (E7335, NEB), have homology to the adapter. The purpose of this step is to introduce barcode sequences and further adapters to enable fragment binding to the sequencing flow cell; the fragments were prepared as before with the following conditions: 3 min 98°C; (15 s 98°C, 30 s 65°C, 30 s 72°C) x20; 1 min 72°C; hold 4°C. The final product was purified again using SPRI beads and then quantified by qPCR following the kit instructions (KAPA Library Quantification Kit, Roche). Fragments were sequenced using an Illumina MiSeq using v3 150 cycle cartridges (MS-102-3001, Illumina), with 5% PhiX loading control (Illumina).

### Data analysis

Sequencing data were processed using a previously described approach ^75^. Briefly, FASTQ data were first demultiplexed by the internal barcodes (introduced by the forward primer of PCR2) using the FASTX-Toolkit (http://hannonlab.cshl.edu/fastx_toolkit/index.html). Samples were then checked for the transposon sequence in 2 steps allowing for a total of 4 bp mismatches. Following identification of transposon-containing reads, the transposon sequence was trimmed and the remaining read mapped using bwa mem (https://bio-bwa.sourceforge.net/bwa.shtml#13), to the *E. coli* BW25113 reference genome (available from the NCBI database, accession CP009273.1, unless otherwise stated). The first nucleotide of a given read was used as the transposon insertion site to generate insertion plot files, which were viewed in Artemis ^76^. Insertion data were cross-referenced with annotated genome features using intersectBed ^77^. We used bbcountunique.sh of BBTools for sub-sampling of FASTQ data (https://sourceforge.net/projects/bbmap/).

### Construction of SPA-tagged strains

A sequential peptide affinity (SPA) tag (containing the calmodulin binding peptide (CBP) and 3×FLAG sequences separated by a TEV Protease cleavage site) was introduced in-frame into the chromosome immediately upstream of the target CDS stop codon, resulting in a C-terminal tag ^53,78^; a tag routinely used for the identification of small proteins ^38,53,79^. The SPA tag was synthesised by GenScript and stitched by PCR ^80^, with a kanamycin resistance cassette flanked by two FRT sites amplified from the *clsA* Keio mutant ^81^. The SPA-Kan^R^ fragment was subcloned into vector pJET1.2 (ThermoFisher Scientific), and the vector used as a template for PCR amplification of the SPA-tag and kanamycin resistance cassette using primers with 50 bp flanking regions homologous to the intended chromosomal insertion site. PCR profile: 98°C 1 min; (98°C 10 s, 62°C 10 s, 72°C 2 min)x34; 72°C 2 min. Gel purified linear fragments were electroporated into BW25114/pKD46 via the Datsenko-Wanner method of chromosomal mutation ^74^. All mutants were verified first by PCR and Sanger sequencing.

### Western blot analysis

Immunoblot assays to determine small protein levels were conducted as described previously with minor modification ^53^. In brief, overnight cultures were sub-cultured 1:500 into LB and grown at 37°C with aeration to either mid-exponential phase (OD_600_ = 0.3-0.6) or stationary phase (OD_600_ = 4.0-5.0). Cells were harvested by centrifugation and resuspended in 50 mM sodium phosphate buffer (pH 8). Re-suspended whole cells were mixed with 4× sample buffer (1× stacking buffer, 2% SDS, 0.025 mg bromophenol blue, 52% glycerol), heated at 95°C for 10 min and centrifuged for 10 min. Samples were separated on Novex 16% Tricine gels (ThermoFisher Scientific) and transferred to nitrocellulose membranes (ThermoFisher Scientific). The membranes were blocked with 2% milk and then probed with anti-FLAG M2-HRP monoclonal antibody (A8592, MilliporeSigma) in PBS-T. Signals were visualized using SuperSignal West Dura Extended Duration Substrate (ThermoFisher Scientific). Equal loading of lanes in each gel was confirmed by examining the relative intensity of background bands in the test samples as compared to the untagged, wild-type control samples.

### Protein structure predictions

Protein secondary structure were predicted using PSIPRED (4.0) with MEMSAT-SVM for TM-domain prediction^82,83^. Protein structures were predicted using ColabFold (v1.5.5): AlphaFold2 using MMseqs2^55^.

### Conservation analysis within E. coli

To investigate the prevalence and distribution of short open reading frames (sORFs) among *Escherichia coli* strains, we screened a collection of globally distributed *E. coli* genomes. We downloaded 227,873 *E. coli* genome assemblies, along with all available metadata, from Enterobase (accessed on 14 March 2024)^84^. The presence of CDSs was determined using tBLASTn (version 2.12.0+). The tBLASTn analysis was conducted with default settings, but adjustments were made to account for the short length of the query sequences: -matrix PAM30 and -word_size 5.

### Identification of small protein homologues outside E. coli

Potential homologues of the *E. coli* small proteins were identified similar to as described ^85^. tblastn searches were conducted of bacteria species genomes using the National Center for Biotechnology Information (NCBI) microbial database. Only species that were labeled as “complete genomes” by NCBI were screened. Each sORF was searched individually. All *Escherichia* (taxid:561) and *Shigella* (taxid:620) genomes were excluded from these searches. Homologues were searched in both “Representative genomes only” and “All genomes – Complete genomes” Organism options. Other settings are as follows: Max target sequences of 1000, Expect threshold of 1000 or 10,000, Word Size of 2, BLOSUM62 Matrix, Gap Costs of “Existence: 11 Extension: 1” and “Conditional compositional score matrix adjustment.” Finally, the low complexity regions filter was turned off in all cases. Unannotated potential homologues were identified through manual analysis of the NCBI Gene database. Potential start codons and stop codons were identified using the Expasy Translate tool to analyze the genomic sequence (web.expasy.org/translate/). Multiple Sequence Alignments were created using the ClustalW program (www.genome.jp/tools-bin/clustalw).

## Supplementary Information

**Supplementary Table 1. Comparison between datasets**

**Supplementary Table 2. Putative protein coding sequences**

**Supplementary Data 1. Protein homolog alignments**

## Supplementary Figures

**Supplementary Figure 1.**
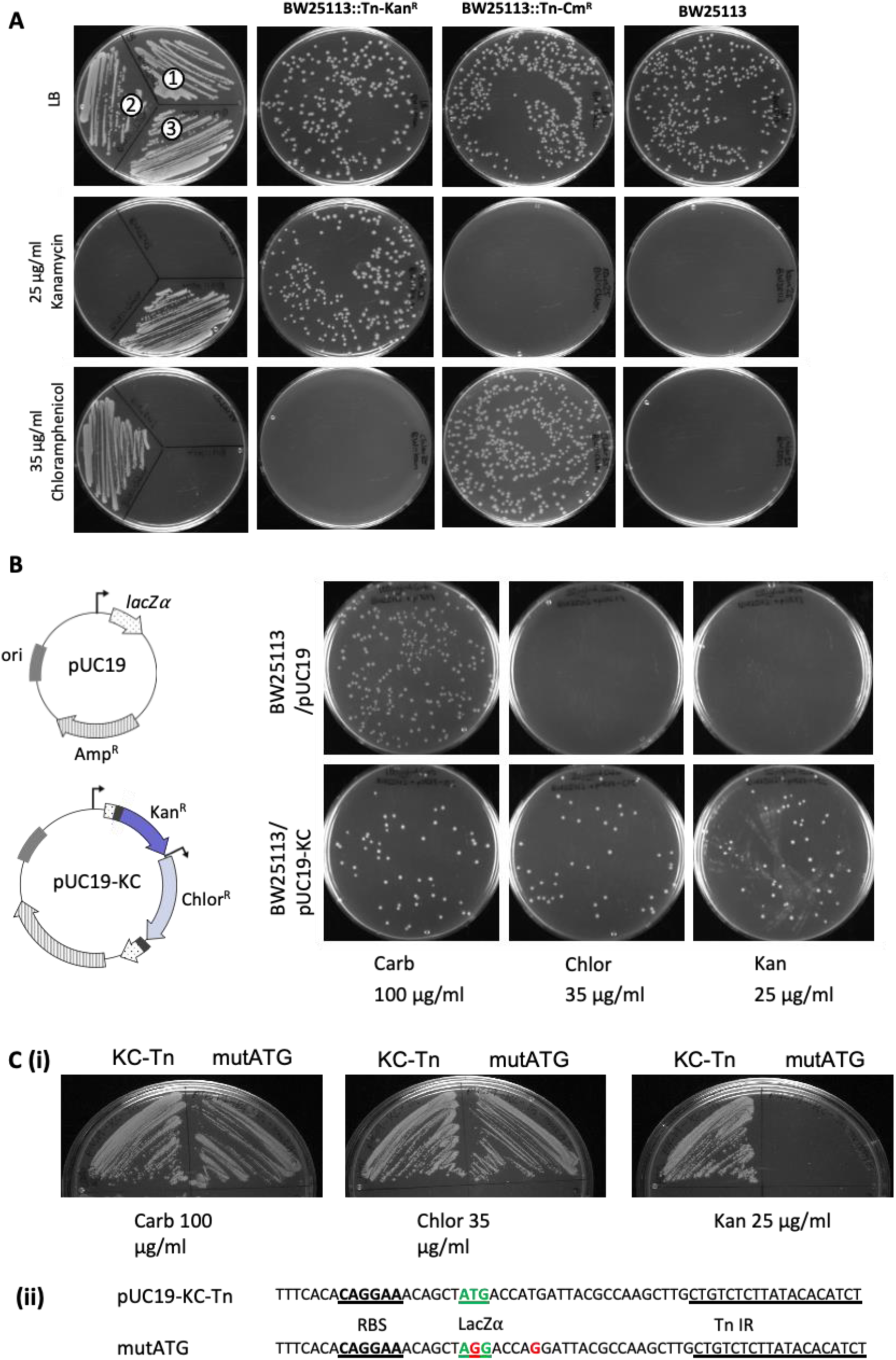
Transposon design and functionality. (A) Antibiotic cassette cross-resistance assay. Growth of *E. coli* BW25113 or *E. coli* BW25113 mutagenized at random with a mini-Tn*5* transposon carrying either a kanamycin (Kan^R^) or chloramphenicol (Cm^R^) resistance cassette on LB agar plates supplemented with antibiotic. The left panel shows the same selection plates with all strains inoculated: (1) *E. coli* BW25113, (2) *E.coli* BW25113::Tn-Cm^R^, (3) *E. coli* BW25113::Tn-Kan^R^. These transposons formed the basis of the dual selection transposon, stitched together by PCR. (B) Translational fusion and expression of the kanamycin resistance gene downstream from and in frame with the LacZα start codon (pUC19-KC). *E. coli* BW25113 was transformed with either the pUC19 plasmid or pUC19-KC on LB supplemented with carbenicillin, chloramphenicol or kanamycin, confirming the LacZα start codon-Tn inverted repeat-Kan^R^ gene fusion is functional. (C) Single nucleotide polymorphisms were introduced by site directed mutagenesis into the start codon(s) of LacZα, replacing ATG with AGG, to make the pUC19-KC-Tn-mutATG vector. *E. coli* BW25113 with either pUC19-KC-Tn or pUC19-KC-Tn-mutATG were inoculated onto LB agar plates supplemented with carbenicillin, chloramphenicol or kanamycin, confirming a start codon is needed for expression of the Tn-Kan^R^ gene.

**Supplementary Figure 2.**
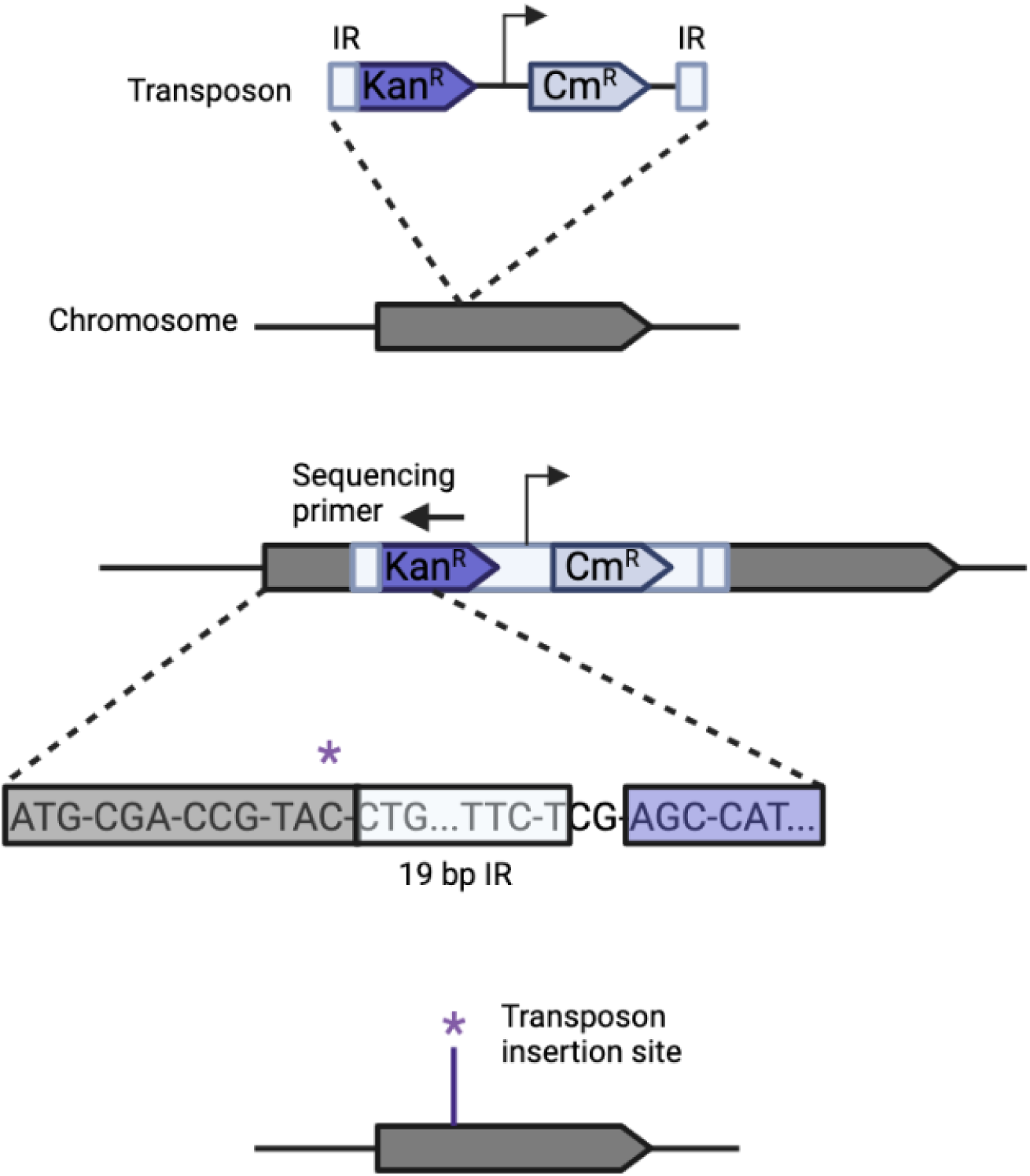
Schematic for identification of the transposon insertion site. The transposon inserts into the genome at random. For identification of the transposon insertion site, the transposon-genomic DNA junction is sequenced by sequencing out from the kanamycin resistance gene into the neighboring genomic DNA. The nucleotide immediately adjacent to the 19 bp inverted repeat of the transposon (indicated by an asterisk) is identified as the transposon insertion site. Abbreviations: IR, inverted repeat; Kan^R^, kanamycin resistance; Cm^R^, chloramphenicol resistance.

**Supplementary Figure 3.**
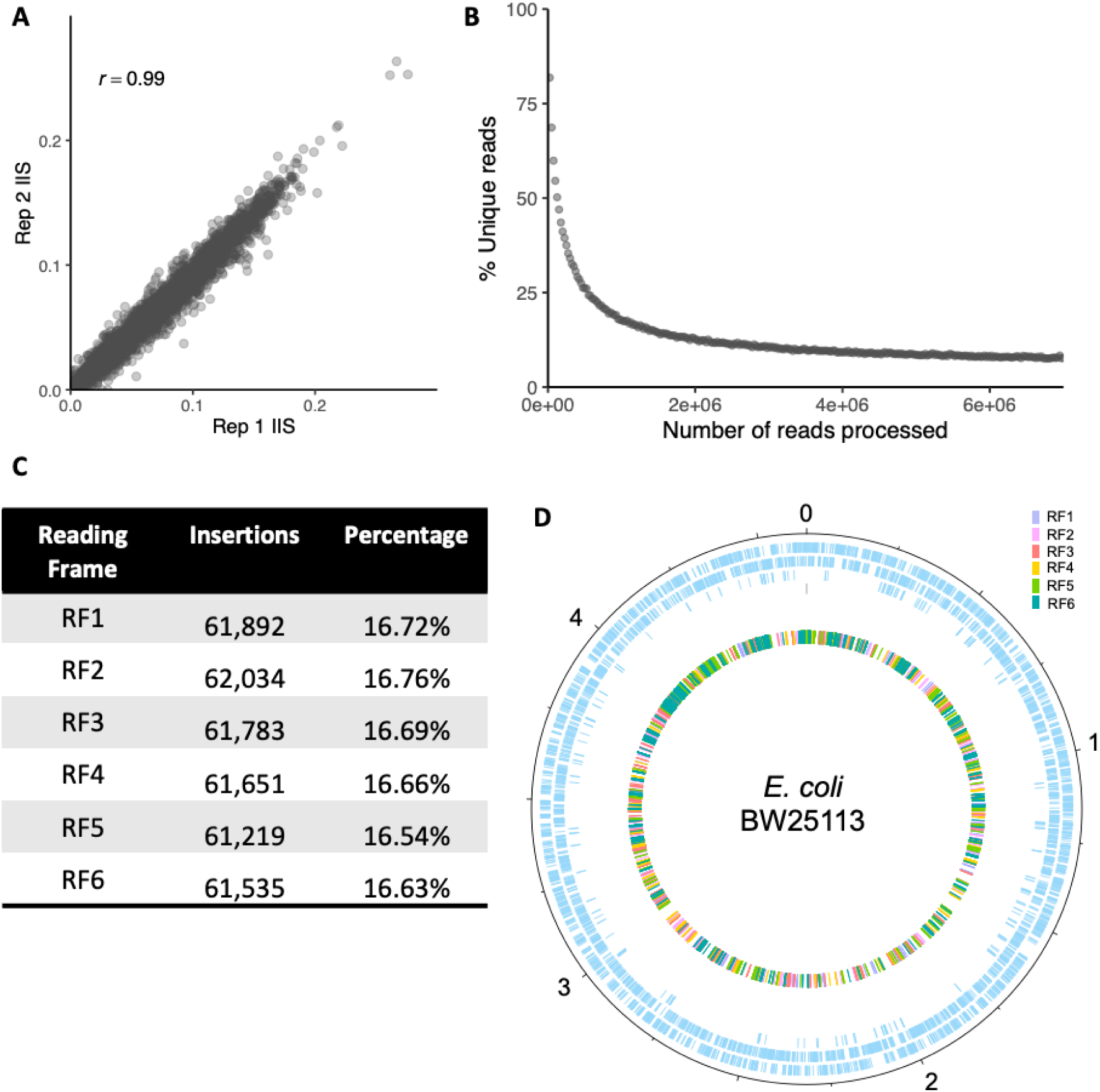
Analysis of the reporter transposon library. (A) Comparison of insertion index scores (IIS; the number of insertions per gene normalised by gene length) between sequenced replicates of the input transposon library. (B) Rarefaction plot of the input transposon library. Sub-sampling of transposon-trimmed fastq data, using BBTools, shows the percentage of unique reads with increasing sampling of sequencing data. (C) Proportion of insertions per reading frame (RF) throughout the genome. (D) Chromosomal position of each identified insertion site around the *E. coli* BW25113 chromosome, and therefore translation reporter position, coloured by reading frame (RF). The two outermost tracks in light blue correspond with sense and antisense CDS respectively, with pseudogenes highlighted in light blue inside.

**Supplementary Figure 4.**
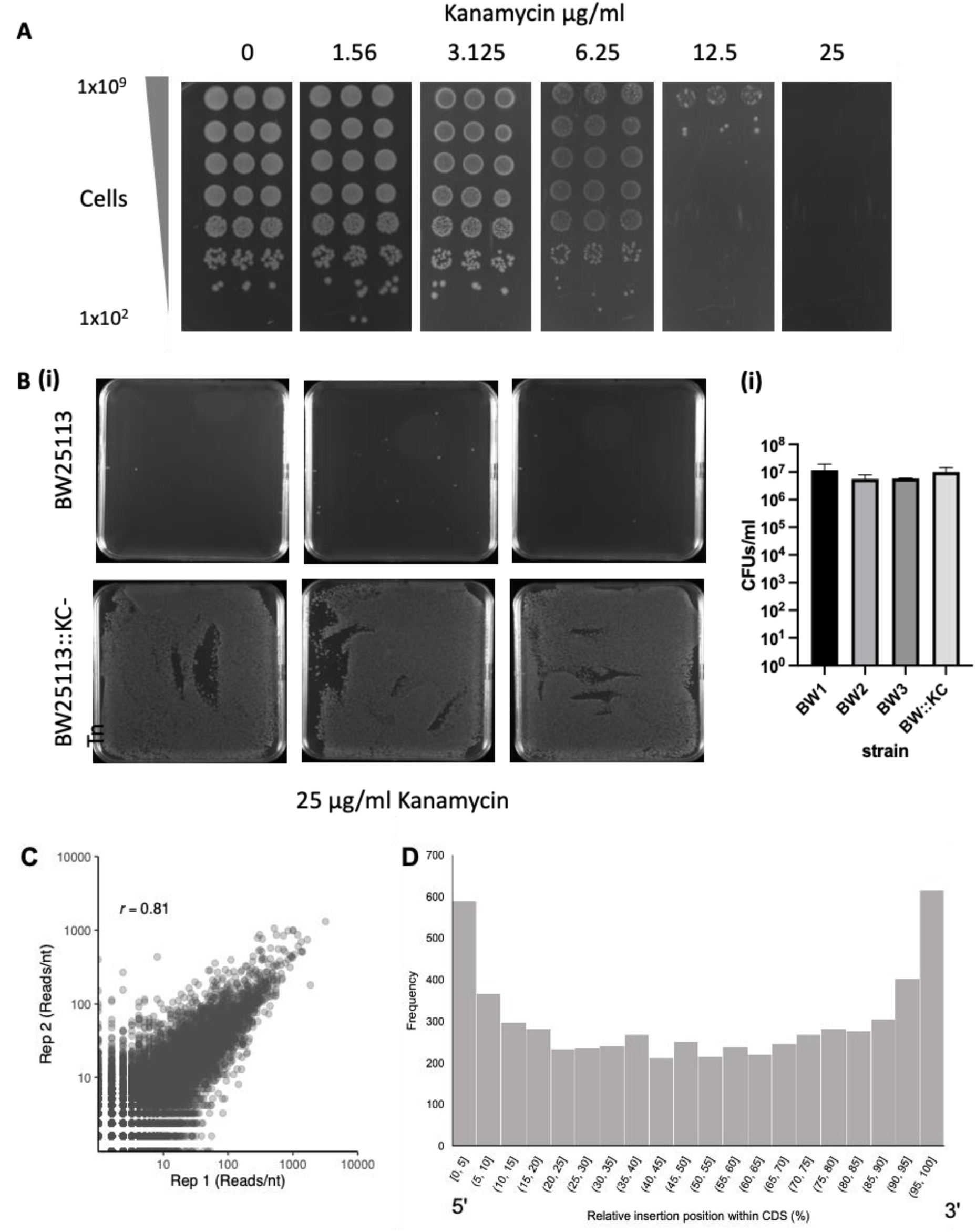
Kanamycin selection screening. (A) Identification of a suitable concentration of kanamycin for agar plate selection. *E. coli* BW25113 was grown in triplicate overnight at 37°C. Samples were normalized to an OD_600_ = 1.00, equivalent to 1×10^9^ cells. Cells were 10-fold serially diluted in LB broth and 5 µl of each dilution was inoculated onto LB agar plates with and without kanamycin at 2-fold decreasing concentrations. Plates were incubated overnight at 37°C and imaged the following day. (B)(i) Cell cultures of *E. coli* BW25113 (in triplicate) and *E. coli* BW25113::kan^R^-Cm^R^-Tn (KC-Tn) were normalized to an OD_600_ = 0.01 in LB; 500 µl of this cell culture was inoculated onto square LB agar plates supplemented with 25 µg/ml kanamycin and grown overnight at 37°C. (ii) In addition, 10-fold serial dilutions of these cultures were inoculated on LB only plates to enumerate the input colony forming units (CFUs) for each sample. (C-D) Analysis of transposon mutants following kanamycin selection. (C) Comparison of the number of reads per nucleotide between replicates of the BW25113::KC-Tn library plated on LB supplemented with kanamycin. (D) The position of each insertion within a protein coding sequence relative to the gene start and end taken as a percentage, following kanamycin selection.

**Supplementary Figure 5.**
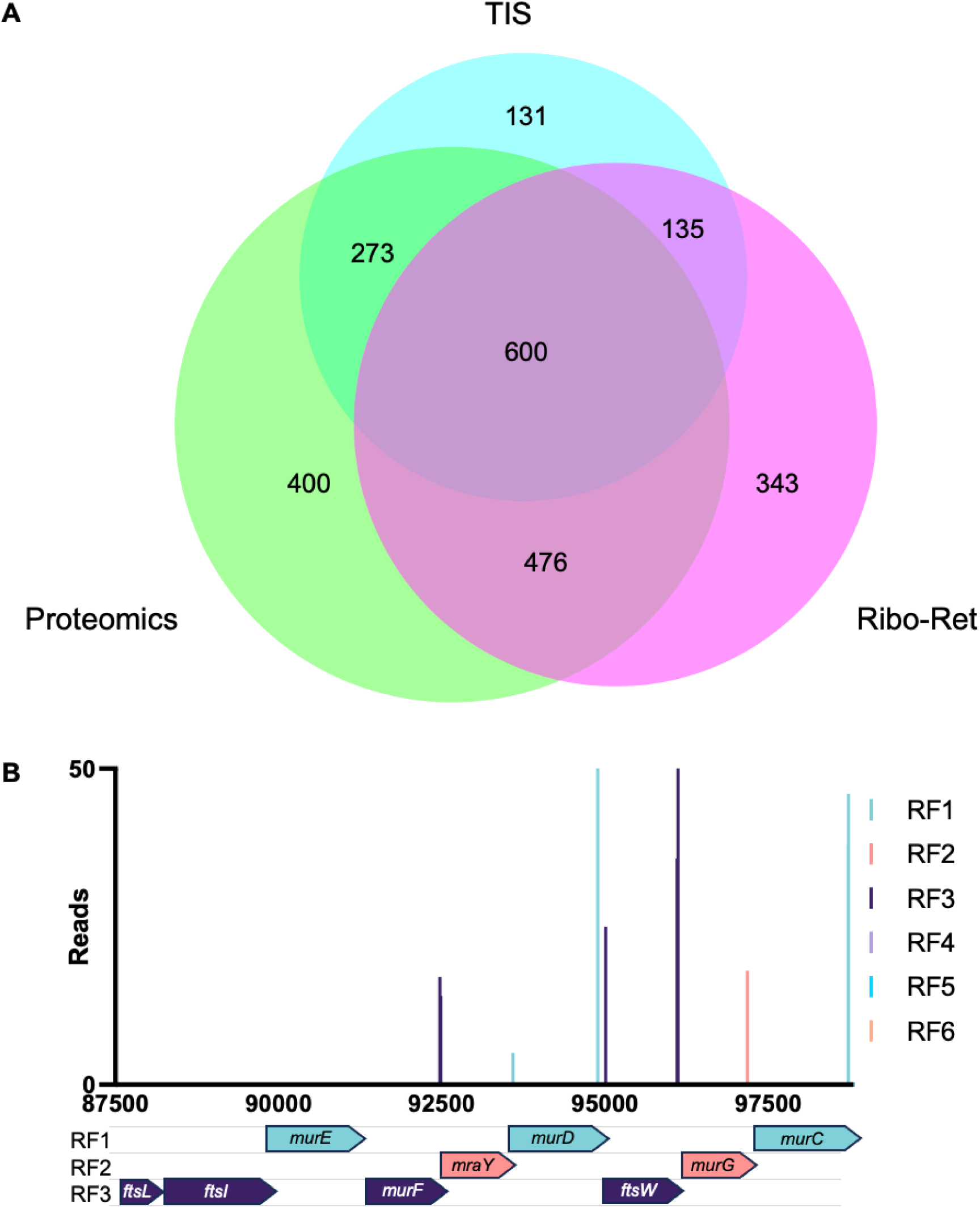
Dataset comparison. (A) Comparison of the annotated genes positively identified by different high-throughput methods for the identification of proteins/protein-coding genes on a whole cell scale. All datasets are reported for the *E. coli* K-12 stain BW25113 grown in LB, with the caveat that the transposon-insertion sequencing (TIS) data was exposed to kanamycin, while the Ribo-Ret dataset was a BW25113 derivative with the genotype BW25113Δ*tolC* and grown in LB supplemented with 0.2% glucose prior to exposure to 12.5 µg/ml Retapamulin. TIS method with a Ribo-Ret dataset (Meydan *et al.* 2019) and a Proteomics dataset (Schmidt *et al.* 2016). (B) Expression of essential genes detected by the translation-reporter transposon. Viable translation-fusion events within 5/9 (*murF, murD, ftsW, murG* and *murC*) essential genes of the *mur* operon following selection of the transposon library on kanamycin. Insertions are at the extreme 5’ end of each CDS, coloured according the reading-frame (RF). The presence of 5’ transposon insertion events suggests translational read-out from the transposon to maintain expression of downstream essential genes.

**Supplementary Figure 6.**
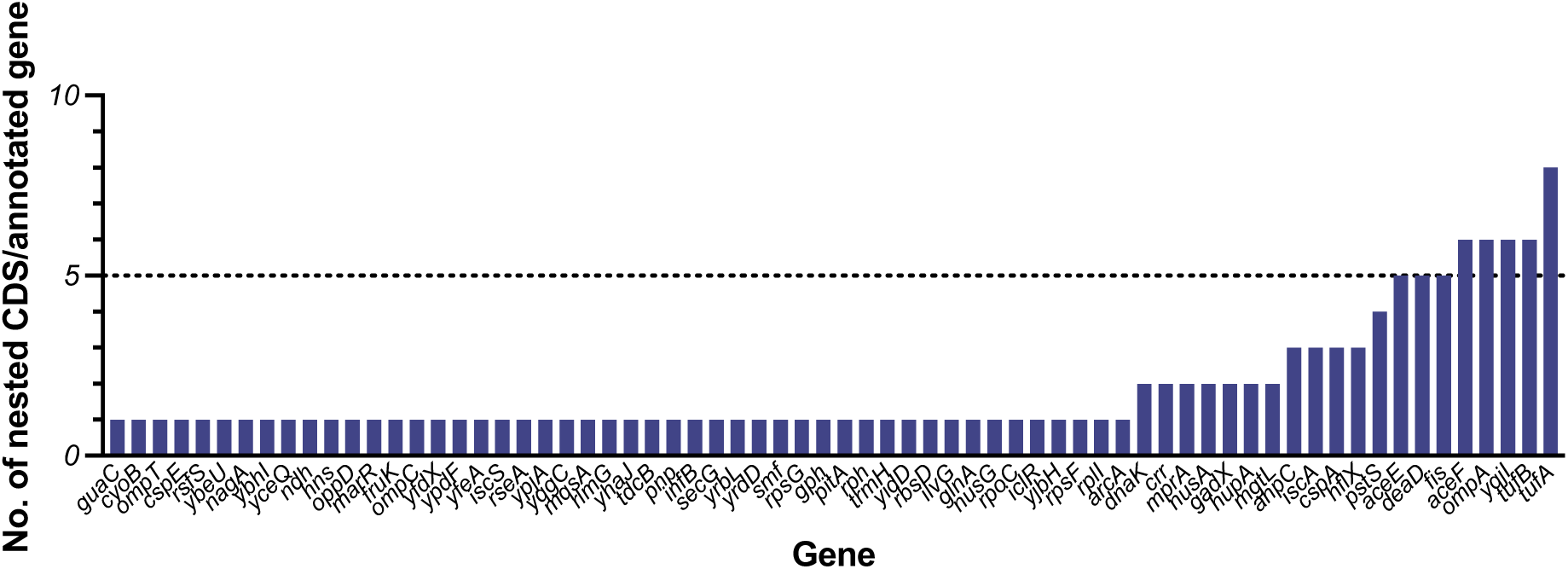
Frequency of out-of-frame nested genes within annotated genes.

**Supplementary Figure 7.**
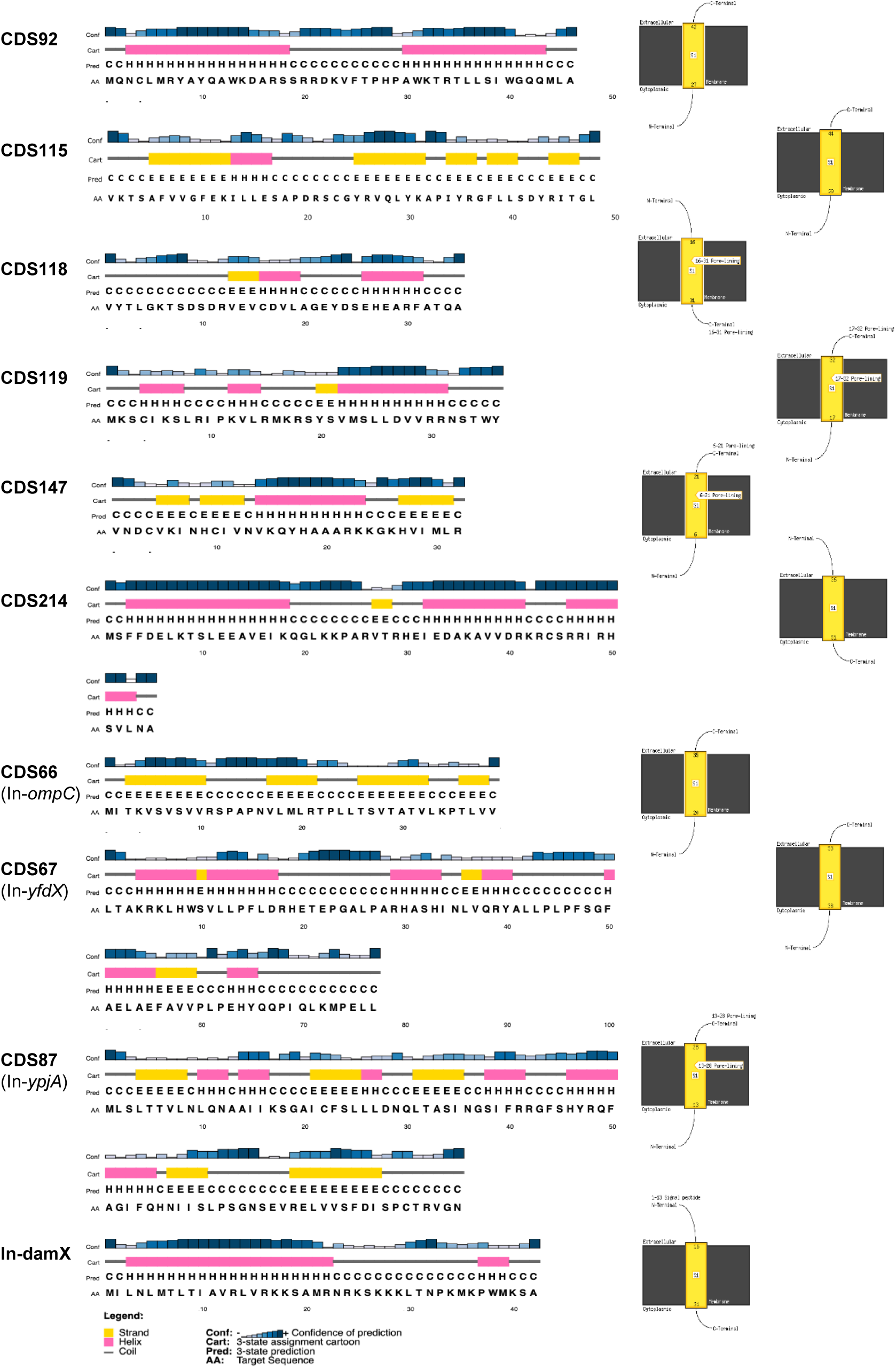
Predicted protein structure using PSIPRED. PSIPRED protein structure predictions, and associated membrane helix prediction using MEMSAT-SVM. CDS42, CDS43, CDS58, CDS61, CDS146, CDS201 and CDS57 were all too short for analysis (minimum size is 30 residues).

**Supplementary Figure 8.**
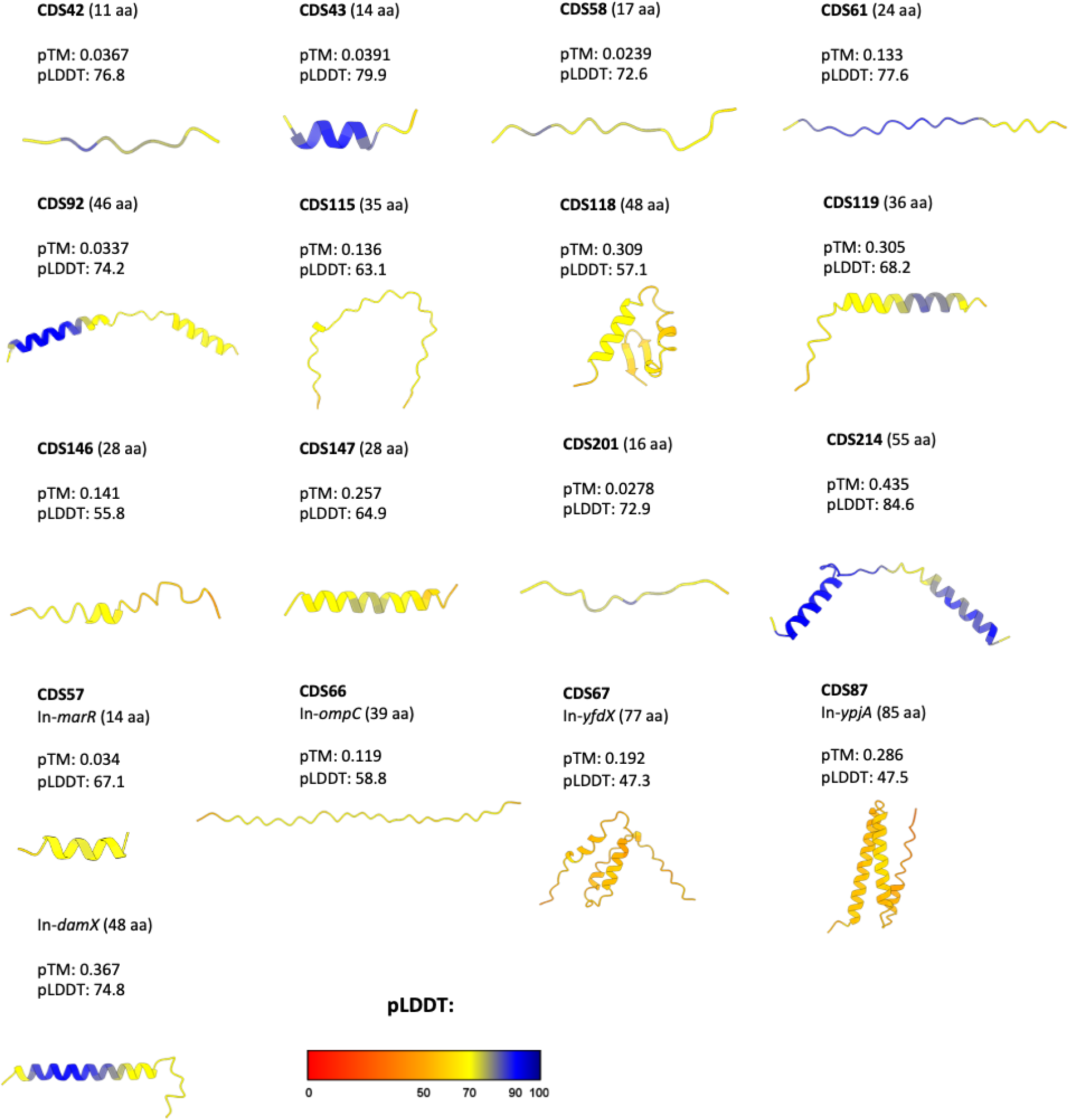
Predicted protein structures. Protein structures predicted using AlphaFold. All structures are shown with the N-terminus on the left and coloured by pLDDT using Chimera.

**Supplementary Figure 9.**
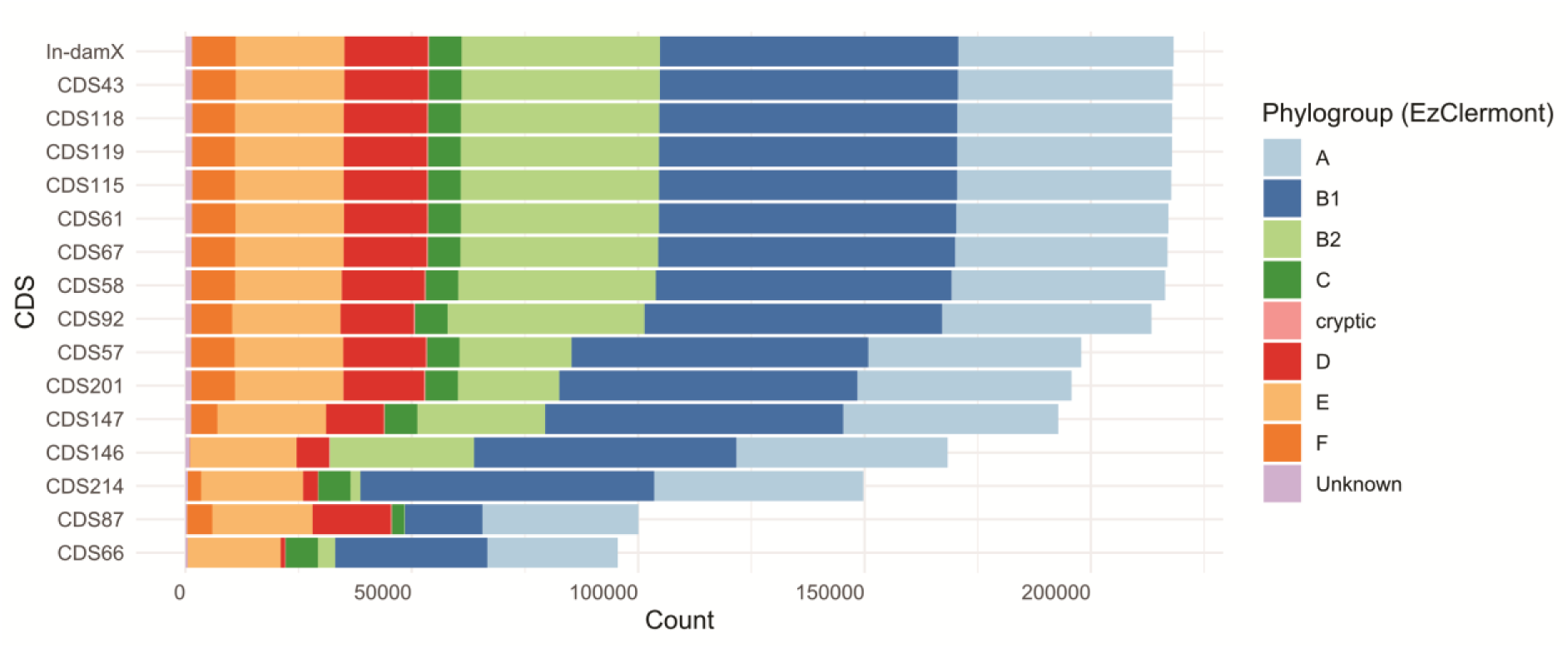
Gene conservation within *E. coli*. The phylogenetic distribution of CDS homologs with ≥80% identity and ≥80% coverage.

## Notes

### Competing Interest Statement

The authors have declared no competing interest.

### Summary of Updates

The Manuscript title has been updated. Additional data regarding conservation of new genes has been added.

